# Effects of weight and height on hand selection: a low-cost virtual reality paradigm

**DOI:** 10.1101/458588

**Authors:** Eric James McDermott, Marc Himmelbach

## Abstract

The main objective was to evaluate the ability of a virtual reality (VR) system to reliably detect the so-called *switch-point* of a user; the distinguishing plane between free-choice use of the left and right hand. Independent variables of height and weight were incorporated into the study design and their effects on hand selection were analyzed. The paradigm utilized the Leap Motion Hand Tracker, along with a custom script written in C# and was realized through a Unity3D application. Stimuli appeared in random locations on the computer screen, and required the participant to reach with the hand of their choice to contact them with a virtual hand inside the virtual space. We observed main effects of height and weight on *switch-points* across the group. We found increased use of the dominant hand as stimuli height increased, as well as a significant increase in overall use of the dominant hand when a 500 gram weighted glove was worn by the non-dominant hand. We validated the average *switch-points* in VR as compared to real-world setups in previous studies. Our results are in line with previously published real-world data, supporting the use of this paradigm in future VR experiments and applications.

## Introduction

Handedness is one of the most obvious asymmetries in the world’s population, with around ~90% identifying as “right-handers” and ~10% as “left-handers” [1]. A range of work has been carried out which suggests that an individual’s selection of the dominant or non-dominant hand is context and task dependent [2–6]. One demonstration of this comes from Gabbard et al [7], who created a manual reaching task where participants were to reach to stimuli at a variety of spatial intervals across 180° space. They found that there was a strong (>95%) preference for the dominant hand at midline and in ipsilateral space. However, this dominant hand preference dropped to under 50% once the targets crossed into contralateral space. This finding suggested that there are competing systems in play: there is a motor dominance factor (i.e. ‘I am right handed’), and there is an attentional system which takes into account the location of the object (i.e. ‘it is in far left space’). They go on to theorize that there must be some of *programming* which takes into account the proximity of the closest effector and the target (*the kinematic hypothesis*), and then the brain takes into calculation the shortest angular distance based on joint and hand space coordinates [8–12]. An alternative hypothesis known as the *hemispheric bias hypothesis* states that each hand is more likely to be used in its ipsilateral space because speed and accuracy are greater in ipsilateral space [5, 13–16]. Furthermore, there is evidence that right-handers also process objects within visual space asymmetrically. That is, right-handers are able to identify stimuli within the right visual-field better than stimuli in the left-half of the visual field [17–18]. Hellige [19] reports 90% of right-handers process visual information in this way, potentially playing a role in the distribution of ‘visually-guided’ reaches. These factors may then altogether enter competition, and resolve in a hand selection and motor movement.

While these studies lay the foundation of hand selection, the concept of task demands has been relatively untouched with regards to effects of height and weight like the current study investigates. In addition, there has been conflicting evidence about the current theories offered. Gabbard [20] did attempt to investigate this concept of task demands further when they blindfolded participants and had them pick up boxes and release them according to auditory cues. They again found that right-handers indeed used their right-hands to reach into ipsilateral space (at a rate of 96%), however, when the task was reversed, the non-dominant hand was chosen only 40% of the time in ipsilateral space. Although these results differed from Gabbard’s earlier experiments, which reported non-dominant hand use closer to 70%, both experiments show an uneven distribution between ipsilateral hand use on the dominant and non-dominant side, providing evidence against the *hemispheric bias hypothesis*. One reason for the increase use of the dominant hand in the latter experiment could be the loss of visual-guided movements, which consequently increases the effort demanded to complete the task.

Building on the work of Gabbard, Przybyla developed a visual feedback / no visual feedback reaching paradigm. They incorporated the use of a horizontally placed mirror to occlude the participants actual hands, and a horizontally placed TV above the mirror to display cursors representing the left and right index finger locations, that were then reflected onto the mirror and seen by the participant. The participants also saw reflections of stimuli targets from the TV monitor on the mirror, and then were tasked with moving one of their non-visible hands to reside as best as possible directly under the target, with or without visual feedback via the help of the cursors. Interestingly, results showed an increase in non-dominant hand use in the no visual feedback conditions; these results are contrary to Gabbard’s findings, and do tend to support the *hemispheric bias hypothesis*. The opposing results perhaps could be due to the difference between the lack of vision and the lack of visual feedback; in one case losing all visual cues, and the in the other losing only the end-effector representation. Apart from this, Pryzbyla had another important finding: subjects used the dominant hand significantly more as target rows moved farther away from the initial starting position. This mimicked results found in an old study by Baldwin [21], who studied his daughter’s reaching preferences in infancy. He found that the farther away an object was while on the midline, the more likely the dominant hand would be chosen.

An array of other task demand experiments have been done. Simon [22] investigated steadiness by measuring movement stability with the preferred and non-preferred hand in a task of placing a small rod through a hole, but found it wasn’t sensitive enough to draw any strong conclusions. Stins [23] had participants pick up different levels of water-filled glasses, but their primary variable of interest in this design was the accuracy needed to pick up the different glasses and the hand trajectories to do so. They did note that different levels of water have different weights, but they did not factor this weight into their analysis and they were only interested in the approach phase before picking up the glasses. Sainburg [24] looked into hand control and trajectory in relation to position, but the targets were always on the horizontal plane. Patla & Rietdyk [25] looked at how obstacle heights influence limb trajectory in locomotion, but nothing regarding so in reaching movements. Cohen and Rosenbaum [26, 27] coined the so-called *grasp-height-effect*, they’ve found an inverse relationship between grasp location on the object and the target position (i.e., you grab something up high on the bottom of the object, and you grab something down low on the top of the object). They’ve also found grasps are done to execute behaviors in the most comfortable positions possible [8, 28–30]. Though Cohen & Rosenbaum collected data on which hand was chosen when objects were in differing locations, they placed an emphasis on the act of grasping the object and putting it in a target location; thereby imposing second-order planning effects of grasp posture that may override the first-order hand selection.

We believe the current work is an original contribution to the field of literature in first-order hand-selection with regard to differing heights, as well as the effects of asymmetrical weight. In essence, we examined the effect that *effort* had on hand selection. We set out to develop a low-cost, quick, reproducible, and robust hand selection paradigm that would accomplish three goals:

1. Investigate *effort-effects* in hand-selection (height & weight), with
2. a hand-selection paradigm developed with state-of-the-art technology in virtual reality that
3. after validation, offers the opportunity to acquire extensive baseline data on hand-selection and create rehabilitation-based tasks with small adaptations to the same experimental setup.

In the last two decades, virtual reality (VR) and virtual reality therapies (VRT) have begun to change the field of neurological rehabilitation and demonstrated their potential in several clinical studies [31–36]. However, clinically certified effective systems often cost over 10,000 euros. To this point, VR has seen crucial advances in accuracy along with considerable reductions to cost in the last 5 years. We propose a paradigm that uses a state-of-the-art virtual reality hand tracker that altogether costs under 100 euros, if the administrator has a minimally viable computer at hand.

Additionally, the flexibility of VR-environments, together with the potential to create highly motivating tasks and procedures, makes VR a potentially powerful tool to measure, diagnose, and rehabilitate motor and cognitive impairments [37]. We propose a paradigm that can make these measurements and accurately record several interesting variables of data in 10 minutes, such as: overall hand-preference, hand-preference by distance, hand-use within ipsilateral space, *effort-effects*, speed of movement, and the trajectory of movement. On top of this, we seek to validate the sensitivity of the paradigm by showing it can distinguish between different grouping conditions (i.e. the *effort-effects*), while maintaining that our findings are in line with prior research results regarding hand-selection, and can be easily adapted to become a rehabilitation paradigm for motor impairments and attentional deficits.

In the present experiment, participants were faced with a virtual environment, in which a grid-like display of cubes appeared in a random sequence one cube at a time. In each trial they were to ‘contact’ the cube with a freely chosen hand and come back to the starting position. The hands were projected into the virtual environment through an infrared hand tracker. If all things were equal, we could expect the use of right and left-hands to fall equally on each side of the midline, however we predict that even in this virtual environment, the results will follow previous studies and show a preference for the dominant hand into near-midline contralateral space. We also expect that the further away a target is from the body’s midline, the more likely the participant will use the ipsilateral hand. Additionally, we predict that increasing the factor of effort will increase the use of the dominant hand in contralateral space. We will test this effort hypothesis in two ways: firstly, by dividing the stimuli presentation into three levels of height; and secondly, in a separate experimental condition, by placing a weighted glove on the non-dominant hand.

## Materials and Methods

### Ethics Statement

The study was approved by the local ethics committee of the University Tübingen and the Medical Faculty Tübingen in accordance with the principles expressed in the Declaration of Helsinki. All participants provided their written informed consent.

### Instrumentation

The experimental setup consisted of the Leap Motion hand tracker (LM) mounted to a desk, a Windows laptop computer meeting the minimum specifications of the LM, and a custom stimuli program created in Unity3D and C#. The LM consists of three infrared cameras and two CCD cameras and integrates custom written algorithms (i.e. Orion Beta) for tracking capability. As stated by the manufacturer, the LM tracks hands and fingers at up to 120 frames-per second and provides a 135-degree field of view with roughly 244 cubic centimeters of interactive 3D space (61cm x 61cm x 61cm). Additionally, the maximum tracking is 80cm from the device, and the sensor’s accuracy in fingertip position detection is approximately 0.01mm. The cost of the LM at the time of submission is approximately $79.99. The laptop computer which ran the LM is an XMG A516 running Windows 10 with a NVIDIA GeForce GTX 1060 graphics card, 16GB of RAM, a solid-state hard drive, and an Intel Core i7-7700HQ CPU 2.80GHz. The cost of this computer at the time of submission was approximately $1399. This computer greatly exceeded the *minimum system requirements* of the LM, which are stated as 2GB of RAM and an Intel Core i3/i5/i7 processor. A computer of these specifications can be found for well under $500.

Grip strength was measured using the Takei Physical Fitness Test “Grip-D” made by Takei Scientific Instruments Co., LTD. A pair of gloves, called “Powergloves”, with pockets to place weighted sandbags were used to impose additional weight to the non-dominant hand. A custom program was written in Unity3D and C# to display stimuli on the laptop screen (www.unity3d.com). Executables and commented source code are available at https://github.com/EricJMcDermott/VR_handselection, under the GPL v3 license. An adapted form of the Edinburgh Handedness Inventory (EHI) [38] was administered to measure the left-hand/right-hand bias of the participants. At the end of the questionnaire, two additional questions were asked regarding the perceived behavioral change in wearing the weighted glove in the latter half of the experiment. The full questionnaire and exemplary responses can be found in the supplementary data (S2 File). Prior to analysis, we discarded the question regarding “with which hand do you use a broom (top hand)” given an overwhelming uncertainty expressed by participants regarding this question.

### Participants

The study consisted of 30 participants (mean age 27.2y, range 19-56y). All participants were healthy, with no signs of or reported medical problems. All were compensated with 5€ at the end of the experiment. Participants must be “right-handed” as defined by scoring higher than the 4^th^ right decile (>74%) on the laterality index in the 13-item augmented EHI. We further excluded participants before data analysis if their overt responses regarding the weighted gloves indicated an explicit strategy in hand selection. Three participants in total were excluded from the data analysis: one participant scored lower than the cutoff for right-handed laterality, one reported that he/she was personally “challenged and motivated to use” the weighted-glove more as if in a gym, and one participant reported explicitly that his job requires extensive ambidextrous hand-use of tools despite a 5^th^ right decile EHI score.

### Experimental Paradigm

The stimuli were generated on the computer screen and composed of 3D represented yellow cubes with shading. In total, the experiment consisted of 1 training round, followed by 4 experimental rounds of stimuli presentation. In the training round, 10 stimuli were presented, whereas in the experimental rounds, 48 stimuli were presented. In both cases, the positions of the stimuli were predefined in a script, and then position presentation was randomized upon initiation of the round. In total, with questionnaires and grip measurements, the experiment lasted approximately 15 minutes.

To begin, participants filled out personal data and consented to the experimental conditions. After which, hand strength was measured using the Grip-D device starting with the right-hand, and then the left-hand. Next, participants were instructed to place their elbows on two corresponding and marked points on the table. Now, the training was initiated and the participant could see their virtual hands on the computer screen. Their hands were seen in a ‘bare-bones’ skeletal view, containing a sphere within the palm region. On the screen, two light-blue squares denoted the region where the participant should center the sphere in the middle of their hand. This represented the starting position (Fig 1).

Once the participants were comfortable in this starting position, they were told about the Leap Motion hand tracker in front of them, and that it functioned best when the participants showed it their palms. Once this was made clear, the participants were told that “yellow cubes will be appearing in the environment one at a time, and you are to reach out and contact the cubes freely with whatever hand that you want to, and the object of the task is to do so as fast as possible.” After the participant finished contacting the first stimulus presented in the training round, they were instructed to come back into the starting position, and repeat this sequence for every stimulus. The next stimulus was presented exactly 2 seconds after the contact to the previous stimulus to provide enough time to return to the starting position while still keeping attention engaged. After the training run of 10 trials, a break screen appeared and the participants were asked if they were ready to go onto the next level, and in this case, the experimental round.

In the experimental condition, the stimuli presentation procedure was the same: the participants started from the starting position, reached out with their chosen hand to contact a cube, returned to the starting position, and then the next cube was presented exactly 2 seconds after contacting the previous one. This was repeated until the break screen came up reporting the end of the round. This break screen consisted of a button that the experimenter would click to initiate the next round, as well as a score that consisted of the summation of centimeters per second that the participant moved in contacting the cubes. This score was explained as the summation of the speed of their movements and the participants were told they would be able to see their scores after every round, but no explicit encouragement nor instruction to improve their score was given.

The stimuli positions in each experimental condition were in a 3-row x 13-column grid. Cube presentations in the middle 3 columns were repeated twice per row in each round, each other column was presented once per row, resulting in 48 trials per round. Each stimulus was a 1×1×1 cube in Unity3D units (which translated to a gain-factor of 2.4 on the computer screen; i.e. 2.4cm × 2.4cm × 2.4cm). The distance between the middle of a cube between two adjacent rows was 1 unit, meaning that the top of a cube in row 1 (the lowest row) would perfectly contact the position of the bottom of a cube in row 2. The distance between columns in Unity3D units was as follows: 1 −1 −.25 −.25 −.25 −.25 −.25 −.25 −.25 −.25 −1 −1, meaning that the outermost 2 cubes on the left and right were bordering each other, while the rest had a 75% overlap with their neighboring cube (Fig 1). Exact grid positions in Unity3D units can be found in supplementary material as (S1 File).

**Fig 1.**
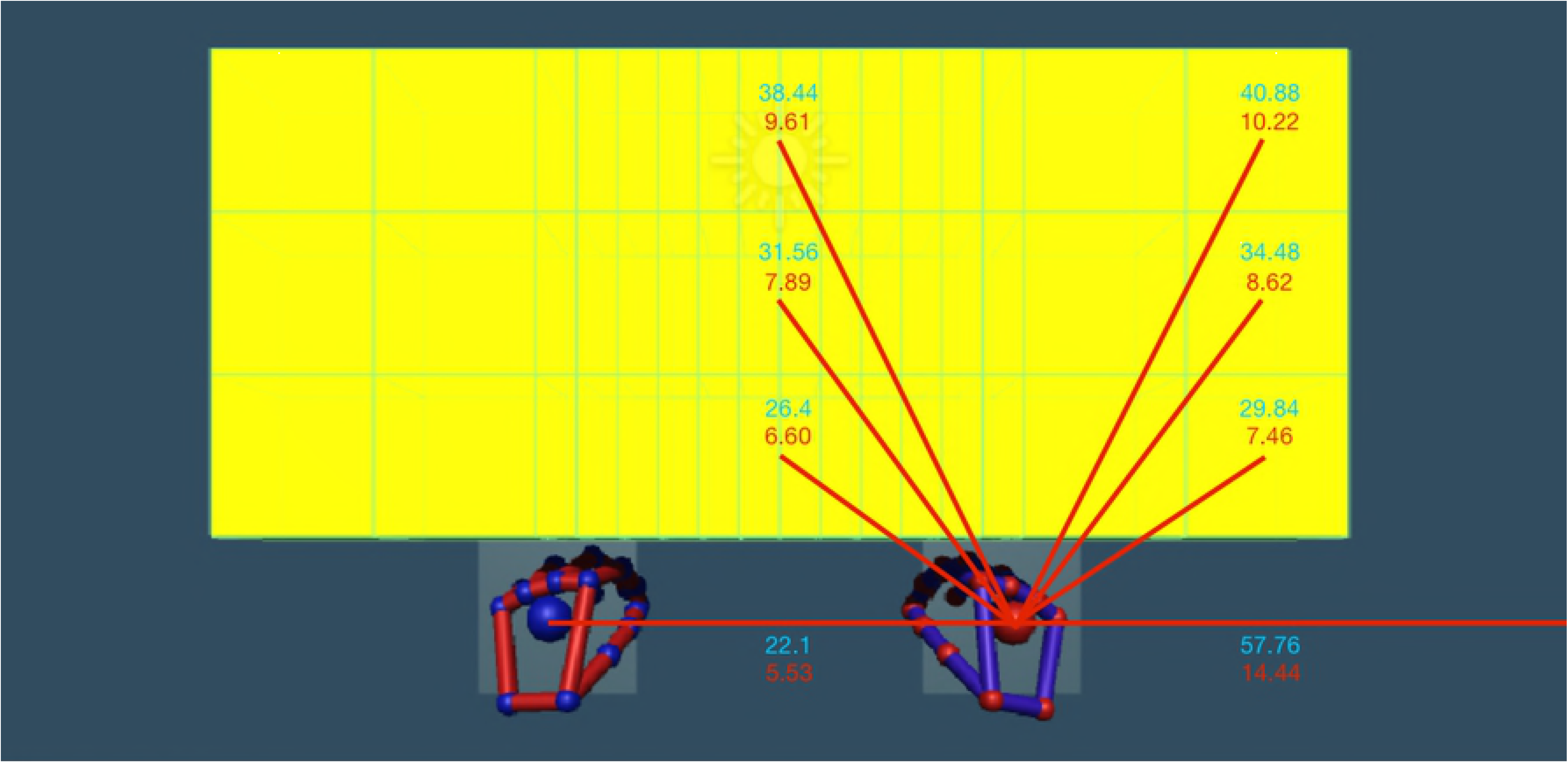
Hand and stimulus configuration. This figure shows all possible stimuli positions on the screen, as well as the initial position of the hands. Each yellow cube is delineated by thin green lines, and was presented independently of the others. The red values are 2D distances moved on the 2D computer screen, and do not account for depth. The black values are the distances the movements required in real 3D space.

Two experimental rounds were conducted consecutively, following which; the participant was instructed to wear two gloves, the right glove contained no additional weight, while the left glove contained a 500-gram sandbag. Visually, they appeared identical. The participant was then instructed again that “you are to reach out and contact the cubes freely with whatever hand that you want to, and the object of the task is to do so as fast as possible.” The participant was asked if they were ready, and then the next round was initiated. Following two experimental rounds with gloves, this portion of the experiment was concluded, and the participant filled out the Edinburgh Handedness Inventory (EHI) and answered complementary questions on their perception of the weighted glove.

### Experimental Setup

The laptop screen measured 19.4cm by 34.4cm. Two light blue squares (2.3cm × 2.3cm on the computer screen) represented the ‘starting positions’ for XY-space, areas in which the respective hand should be centered within at the start of each trial. Centered in the starting positions, the hands were presented 5.50cm away from each other, and 14.45cm from the left and right edges of the screen (Fig 1). The actual starting positions of the participants hands were in a different Z-plane in depth from the target stimuli, an average of 5.37cm away from the target stimulus plane due to the slight variations in arm length. Stimulus positions on the screen as well as measured kinematic data from the LM tracker were all in arbitrary Unity3D coordinates and were analyzed as such (please see below ‘Data Collection and Statistical Analysis’). We later converted results to visual and real space coordinates to allow for a better understanding, interpretation, and direct comparisons with previous studies. In the following, we report distances and positions in the experimental setup in two reference frames, visual 2D and visual 3D space. First, in correspondence with the 2D visual space of the laptop screen to describe the actually visible stimulus movements on the screen (visual2D). Second, in correspondence with the virtual 3D space taking into account the simulated visual depth of hand and target representations on the screen (visual3D). The distance from the hand starting position of either hand to the center of the box in the bottom row 1 of the middle column was 4.2cm in visual2D, and 6.60cm in visual3D (Fig 1 and Fig 2); reaching distance to the center of the box in the middle row 2 of the middle column was 5.8cm in visual2D, and 7.89cm in visual3D; reaching distance to the box in the top row 3 of the middle column was 7.9cm in visual2D, and 9.61cm in visual3D. The distance to reach to the middle of the box in row 1 of the most peripheral column was 4.6cm in visual2D and 7.46cm in visual3D; reaching distance to the box in row 2 of the most peripheral column was 6.3cm in visual2D and 8.62cm in visual3D; while the distance for the box in row 3 of the most peripheral column was 8.5cm in visual2D and 10.22cm in visual3D.

In a third reference frame, real-space, the hands were held 22.1cm away from each other in correspondence with the starting positions presented on the screen. The elbows were placed at markers on the table exactly 30cm from each other (Fig 2). Each participant was instructed to maintain an arm angle that resulted in a comfortable position, while also extending forward from the elbows and placing the hands slightly inwards so that the center of the virtual palm resided in the starting positions. This position was visually confirmed by the experimenter, who was present at all times. To calculate real 3D reaching distances we used a rounded gain factor of 4, which was found by dividing the real measured distance the hands were held apart in order that the hands were in the starting positions (22.1cm) and the average distance the visual representations of the hands were held apart on the computer screen (5.53cm). Therefore, the calculated real space distance to the center of the boxes within the middle column of row 1 was 26.4cm; row 2, 31.56cm; and row 3, 38.44cm. (Please see Fig 1 and Fig 2 for further measures.) The LM tracker was positioned 34.8cm diagonally from the bottom corners of the computer screen, and 40cm diagonally from each elbow marker (Fig 2).

**Fig 2.**
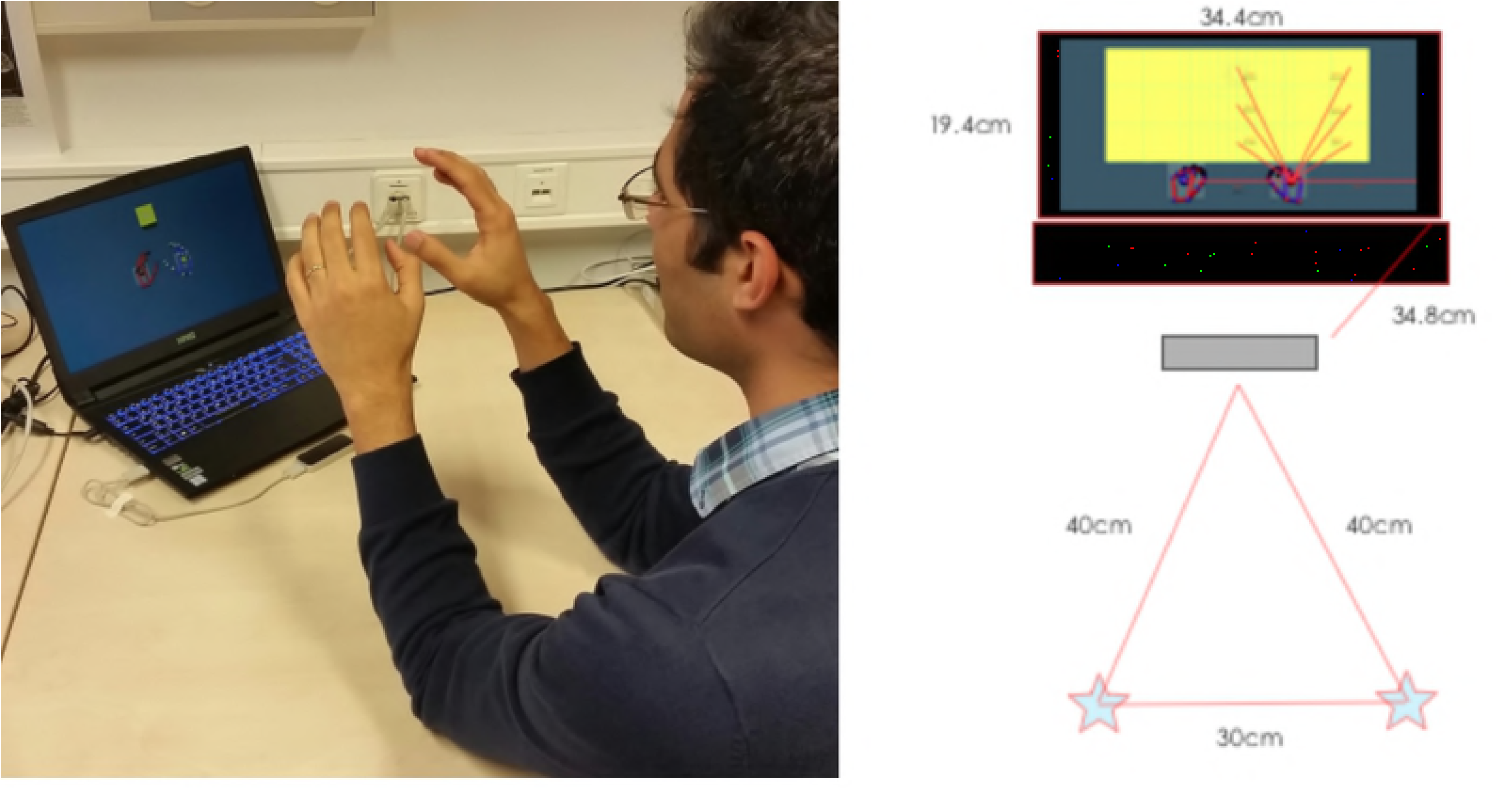
Experimental setup. The left panel shows a participant in the “initial starting position” before going to contact the cube. The right panel shows the same setup in a schematic drawing with the distances on the setup. Respectively, the stars are where the elbows were placed, the grey rectangle is the Leap Motion, and the black rectangles are the keyboard and computer monitor.

### Data Collection and Statistical Analysis

Kinematic data was collected in real-time during the entire duration of the experiment using the LM tracker and custom-written Unity3D scripts. Information that was collected included: 3DCoordinates of both hands upon stimulus presentation, timestamp upon stimulus presentation, coordinates of stimulus presentation, 3D trajectory of the hands until contacting cube at a 20 Hz sampling rate, timestamp upon hand contact with stimulus, and which hand contacted the stimulus. Additionally, we recorded EHI scores, grip strength of both hands, and questions regarding the experience wearing the weighted glove.

Data was used from the experimental rounds only, the training round was not included in the analyses. Within each row, condition, and participant, we sorted stimuli according if they were touched with the left or the right hand. The horizontal distance from midline was averaged across all cubes contacted with either the right or the left hand, resulting in left hand and right hand horizontal mean positions. The sum of these means was first divided by 2, then by the total range of horizontal stimulus positions within the experiment; these set divisional values were the same for each participant, condition, and row. Finally, the resulting sum was multiplied by 100. The resulting number estimated the virtual point at which hand-use switched from one hand to the other on a continuous scale, i.e. the *switch-point* within each row. A *switch-point* was determined for each participant at each height, i.e. row of stimuli, and for each condition, i.e. weighted and non-weighted.

We then analyzed switch points with a 3 × 2 factorial repeated-measures ANOVA for the main effects of height (bottom, middle, & top row) and weight (non-weighted vs. weighted), and the interaction. Significant findings for the main effect of height or the interaction of weight and height were followed up with subsequent rANOVAs and paired t-Tests. Bonferroni-corrections for multiple comparison have been applied to the respective post-hoc tests. Assumptions of sphericity have been tested with Mauchly-tests and Greenhouse-Geisser corrections have been applied to DFs in case of a significant violation (Mauchly-tests p < .05).

Please note, that all measurements, calculations, and analyses were conducted with Unity3D coordinates. In doing so, we avoided any variability or systematic biases from rounding errors. Only final group results were converted to the three reference spaces, i.e. visual2D, visual3D, and real space.

## Results

The rANOVA showed a significant main effect for non-weighted (NW) versus weighted (W) conditions (p = 0.009, F_1,26_ = 7.9582; Tab 2 and Fig 3), with a mean difference (NW - W) of −0.64cm, SEM 0.06; i.e. average switch points moved significantly towards the left from the non-weighted to the weighted conditions. We also found a significant main effect of height (p = 0.0107, F_2,52_ = 4.960). We followed this up with three paired t-tests comparing switch points between height levels. These analyses revealed a significant difference between the bottom vs middle row (mean difference bottom - middle: 0.64cm, SEM: 0.064, p = 0.0006) and the bottom vs high row (mean difference bottom - high: 1.04cm, SEM: 0.112, p = 0.0305), with a non-significant result for the comparison of middle vs high (mean difference middle - high: 0.4cm, SEM: .104, p = 0.7932). Yet, after correcting for multiple comparisons using a Bonferroni-corrected error probability threshold of α = 0.05/3 = 0.0167; only the bottom vs middle row stayed significant at the threshold. The interaction of the factors height and weight was not significant (p = 0.3528, F_2,52_ = 1.0630).

**Table 1.** Mean switch points and standard errors of the mean (SEM) in real space, in visual2D space, and in Unity3D units.

|  | Non-Weighted | Weighted | Mean (SEM) |
| --- | --- | --- | --- |
| **Top row** | -2.04cm (0.492)<br>-0.51cm (0.123)<br>-0.2125 (0.0514) | -2.72cm (0.532)<br>-0.68cm (0.133)<br>-0.2838 (0.0553) | -2.38cm (0.464)<br>-0.594cm (0.116)<br>-0.2481 (0.0483) |
| **Middle row** | -2.16cm (0.348)<br>-0.54cm (0.087)<br>-0.2257 (0.0362) | -2.48cm (0.444)<br>-0.62cm (0.111)<br>-0.2568 (0.0462) | -2.32cm (0.360)<br>-0.58cm (0.090)<br>-0.2413 (0.0376) |
| **Bottom row** | -1.20cm (0.296)<br>-0.30cm (0.074)<br>-0.1240 (0.0307) | -2.16cm (0.352)<br>-0.54cm (0.088)<br>-0.2255 (0.0368) | -1.68cm (0.296)<br>-0.42cm (0.074)<br>-0.1748 (0.0309) |
| **Mean (SEM)** | -1.80cm (0.340)<br>-0.45cm (0.085)<br>-0.1874 (0.0353) | -2.44cm (0.4)<br>-0.61cm (0.1)<br>-0.2554 (0.0416) | -2.04cm (0.352)<br>-0.51cm (0.088)<br>-0.2214 (0.0367) |

**Fig 3.**
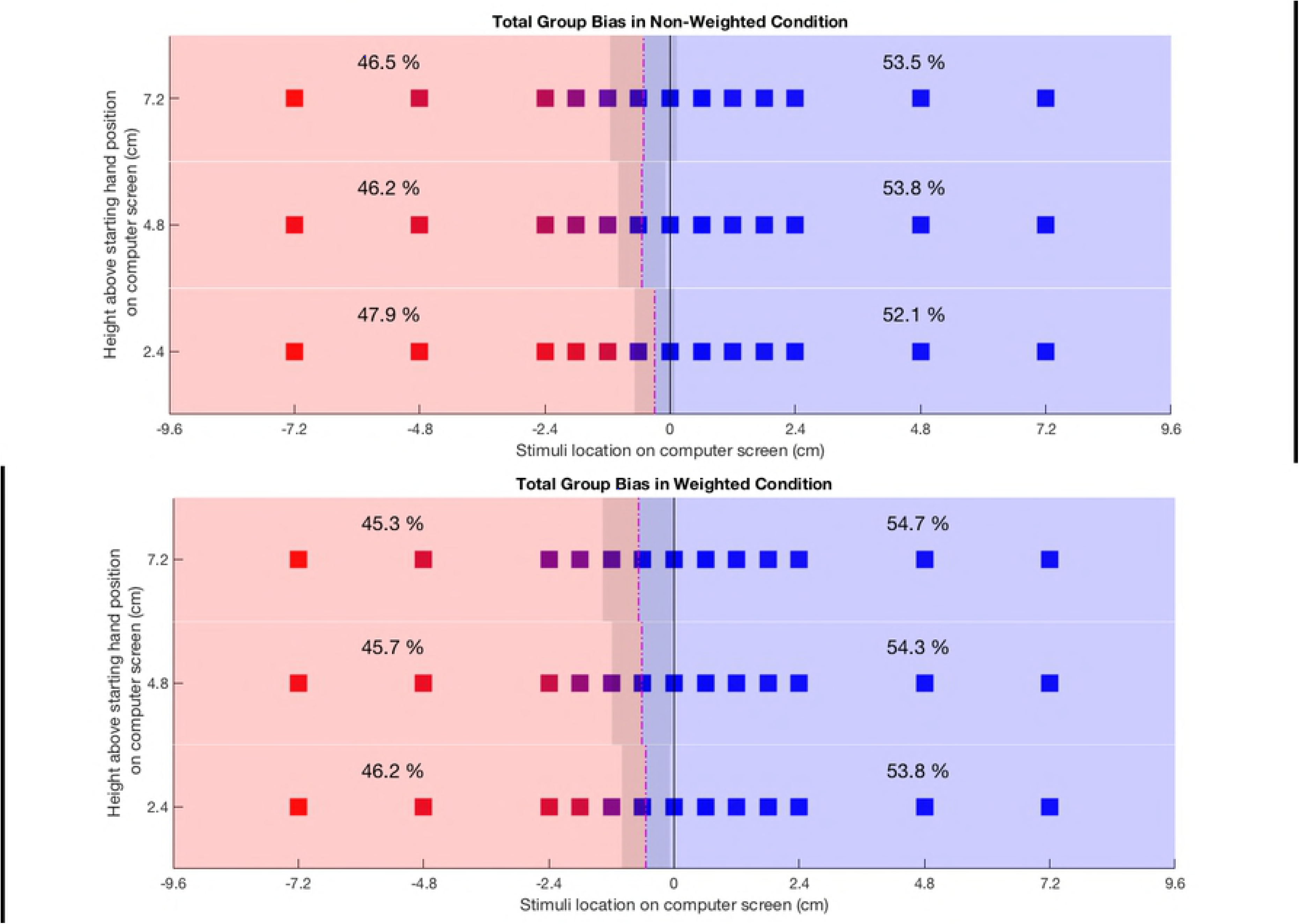
Average switch points. Group means of hand selection at stimulus positions, group mean switch points, and mean percentages of left and right hand use, respectively. Each cube represents a stimulus location, pure blue corresponds to stimuli contacted only with the right hand, while pure red represents those contacted only with the left hand. Dash-dot lines indicate mean switch points with shaded areas representing the standard deviation of switch points. Upper panel: non-weighted conditions; lower panel: condition with weighted left hand glove.

Beyond this main analysis, we conducted complementary analyses to explore the participants’ behavior and compare our data to previously reported findings. We calculated the number of midline crossings for either hand. We found a group mean of 9.89 crossovers in the NW-condition that increased to a 12.11 crossovers in the W-condition for the right hand. In contrast, the left hand went from 3 crossovers in NW-condition to 2.59 crossovers in the W-condition. We calculated the frequency of ipsilateral hand use in ipsilateral space. This data showed that the participants used the dominant right-hand on average 93% of the time in ipsilateral space, whereas the non-dominant left hand was used on average 76% in ipsilateral space (Fig 4). Beyond this difference, variability across the group was apparently higher for the non-dominant left hand than for ipsilateral right hand use.

**Fig 4.**
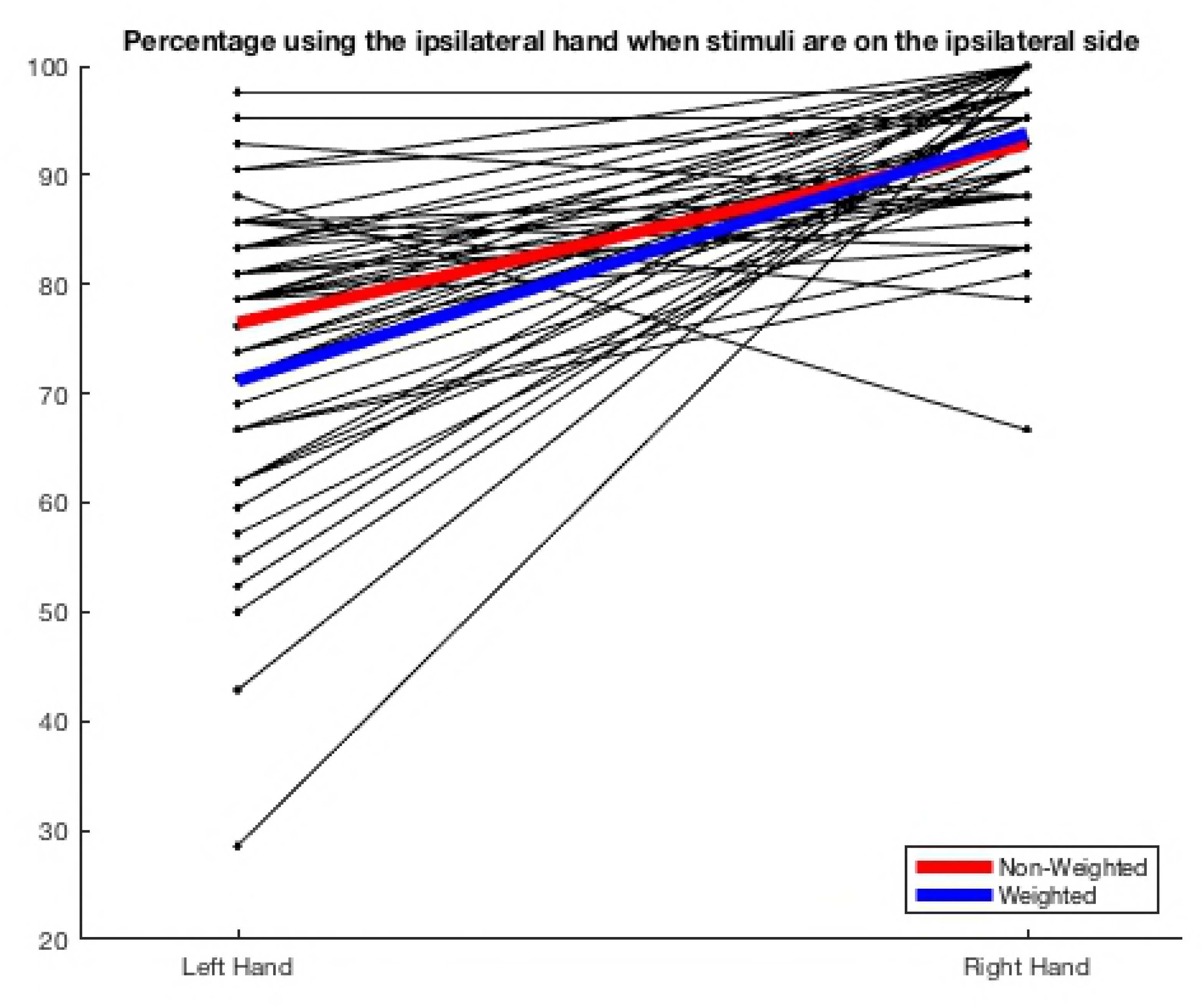
Ipsilateral hand selection. Mean percentage of spatially congruent hand selection, i.e. selection of left and right hand use, respectively, for ipsilateral targets (thick red and blue lines). Individual values from each participant are shown as thin, black lines.

To determine if all participants complied with the instructions and started in the same locations, we analyzed starting positions across the group. We read out horizontal and vertical coordinates of both hands at the time of target presentation for each trial and each participant. From this data we calculated frequencies of positions within each participant and then averaged these frequencies across the group (Fig 5). This analysis showed that a highly consistent percent of trials began within the same square centimeter of space, with the left hand having a mean horizontal starting position of −2.9cm, and the right hand having a mean horizontal starting position of +2.9cm.

**Fig 5.**
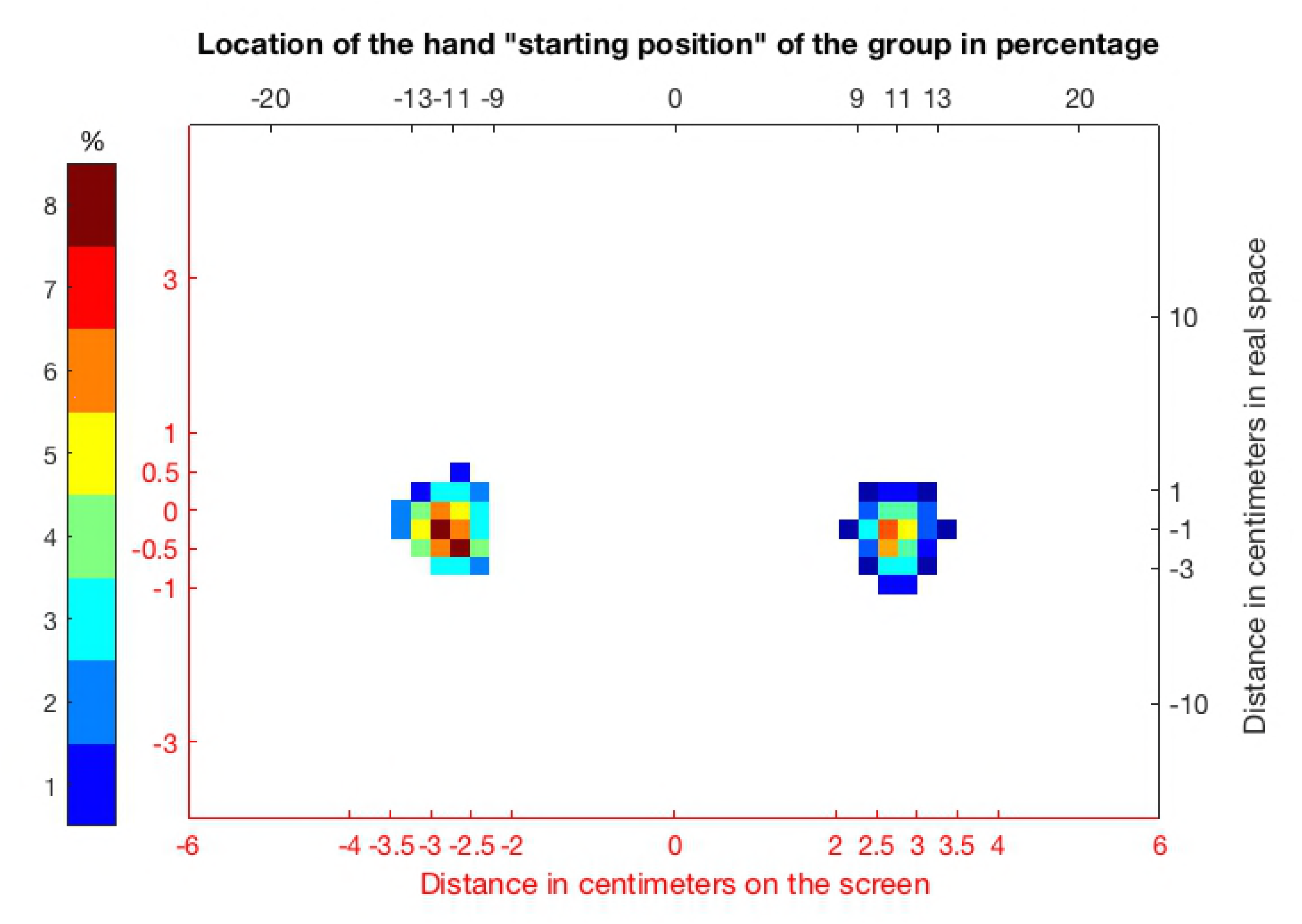
Spatial distributions of hand starting positions. Data indicate the group mean percentage of occurrence of a given position within the individuals’ number of trials. Red axes indicate positions in centimeters on the screen (visual2D), while black axes indicate positions in real space (please rf. to Fig 1 and Fig 2).

We also collected data from the Edinburgh Handedness Inventory (EHI). The average score on the EHI was 92.02, with a standard deviation of 8.6 and a range of 80.77 – 100. This places our average participant in the 8^th^ decile for right-handedness. We correlated handedness score, switch-point measures (switch-points for each weight condition at each height level and their respective difference values between weighted and non-weighted condition), and grip strength data (grip strength left, grip strength right, difference between grip strengths) across our group of participants. This correlation analysis comprised 13 variables and thus resulted in 78 correlation coefficients. The Bonferroni-corrected threshold for these analyses was α = 0.05/78 = 0.000641. Only correlations within groups of measurements, i.e. between switch points from different conditions and/or rows, survived this threshold (Tab 3). If we also took uncorrected findings into account, we saw two correlations between grip strength measures and switch points (grip strength left and switch points weighted condition, high row; grip strength difference and switch point difference between conditions, middle row; Tab 3). With a negative sign, both correlations indicate an expected direction of associations, the stronger the left hand (relative to the right hand), the more rightward the position of the (change of the) switch points. Even at uncorrected thresholds we found no correlation of EHI scores, neither with grip strength measures nor with any switch measures.

**Table 2.** Correlations between dependent variables of hand use.

|  | **gripR** | **switch**<br>**NWM** | **switch**<br>**NWH** | **switch**<br>**WL** | **switch**<br>**WM** | **switch**<br>**WH** | **switch**<br>**DiffL** | **switch**<br>**DiffM** | **switch**<br>**DiffH** |
| --- | --- | --- | --- | --- | --- | --- | --- | --- | --- |
| **gripL** | **0.926**<br>**4\*10<sup>-12</sup>** |  |  |  |  | -0.41<br>0.0322 |  |  |  |
| **gripD** |  |  |  |  |  |  |  | -0.39<br>0.0472 |  |
| **switch**<br>**NWL** |  | **0.73**<br>**1\*10<sup>-5</sup>** | **0.64**<br>**0.0003** | **0.68**<br>**0.0001** | **0.74**<br>**1\*10<sup>-5</sup>** | 0.57<br>0.0020 |  |  |  |
| **switch**<br>**NWM** |  | - | **0.71**<br>**3\*10<sup>-5</sup>** | **0.66**<br>**0.0002** | **0.66**<br>**0.0002** | **0.68**<br>**0.0001** |  |  |  |
| **switch**<br>**NWH** |  |  | - | **0.67**<br>**0.0001** | **0.68**<br>**9\*10<sup>-5</sup>** | **0.64**<br>**0.0004** |  |  |  |
| **switch**<br>**WL** |  |  |  | - | **0.83**<br>**6\*10<sup>-8</sup>** | **0.63**<br>**0.0004** | -0.58<br>0.0016 | -0.41<br>0.0327 |  |
| **switch**<br>**WM** |  |  |  |  | - | **0.72**<br>**2\*10<sup>-5</sup>** |  | **-0.63**<br>**0.0004** |  |
| **switch**<br>**WH** |  |  |  |  |  | - |  |  | -0.49<br>0.0087 |

Please note that only correlations with $p < 0.05$ uncorr. are presented. Significant findings after
Bonferroni-correction are shown in bold. gripL: grip strength left; gripR: grip strength right; gripD:
grip strength difference; NWL: non-weighted low; NWM: non-weighted middle; NWH: non-
weighted high; WL: weighted low; WM: weighted middle; WH: weighted high; DiffL: switch point
difference low; DiffM: switch point difference middle; DiffH: switch point difference high.

## Discussion

The finding of a main effect of weight supported our hypothesis that an increased amount of effort would shift the switch-point farther into the contralateral space of the dominant hand. To our knowledge, this is the first study which incorporates asymmetrical weight as an independent variable. We offer the *effort-effect* hypothesis to explain the results: given the extra effort needed to use the non-dominant hand in this condition, the selection factor for the dominant hand “took over” on several of the contralateral targets. We define our *effort-effect* hypothesis as follows: “given an initial preference to the dominant hand at midline, we expect thereafter that the more effort it takes to use a hand in comparison to the other, the less likely it will be selected for use due to the increased cost-of-movement. If increased effort would be required equally between both hands, in the case of increased distance on the midline equal between the two hands, preference will be increased for the dominant hand.”

The main effect of height also supports this *effort-effect hypothesis*, in that the middle and top rows showed significantly more dominant hand use compared with the lowest row, where the stimuli were closest to the hand starting position. This evidence also directly corresponds with the findings by the Baldwin [21], who administered over 2000 reach preference trials to his daughter from her 5^th^ month to her 10^th^ month. Baldwin noticed that at a distance of 9in (~23cm), his daughter showed no trace of hand preference. Yet, when he increased the distances to 12-15in (~30-38cm), he was able to record a preference ratio of 15:1 for the dominant hand. Baldwin concluded that this selection preference was due to the extra exertion of effort that the farther targets demanded. This concept also follows Fitts’ Law [39], that the modulation of effort from task difficulty should indeed increase with target distance. Interestingly, Pryzbyla [6] reported a finding with a different stimulus setup which also is complementary with our hypothesis. The targets in their experiment required participants to reach farther *in-depth* rather than height, but despite that, they found a similar effect that the dominant right hand tended to be used increasingly more with increasing depth.

Additionally, we calculated the frequency of ipsilateral hand use in ipsilateral space. In the non-weighted condition, the data showed that the participants used the dominant right-hand on average 93% of the time in ipsilateral space, whereas the non-dominant left hand was used on average 76% in ipsilateral space (Fig 4). Being such, it also follows that this paradigm is representative of the literature in distinguishing hand selection. In particular, a study by Harris and Carlson [40] found that the dominant right hand was used 90% of the time with ipsilateral stimuli, where the non-dominant left hand was used 60-75% of the time with ipsilateral stimuli. This agreement with past literature is also in itself significant, as in our knowledge, this is the first virtual reality paradigm to address hand selection in this way. In line with the *hemispheric bias hypothesis*, we found a large percentage of ipsilateral reaches, possibly driven by reaction-time advantages due to spatial compatibility between the arm and target [16]. The smaller percentage of contralateral reaches could be explained by the *kinematic hypothesis*, in that occasionally the joint angles lend themselves to make more direct paths to a target in contralateral space. This also follow the logic of Przybyla [6], in that “one should not always be expected to use one arm, but rather individuals should tend to choose the arm that is more proficient for the task conditions at hand.” Yet, the *kinematic hypothesis* would not offer a solution when we bring the idea of task demands regarding weight or height into the picture. However, these findings can be resolved by the addition of our *effort-effect hypothesis*, in that the initial preference of the right-hand takes precedence on some of the stimuli locations that are in contralateral space, and then even more so when the left hand is ladened with the 500g weight. To support this, we looked at the number of crossovers at each position. In total, there are the 267 right-handed crossovers in the group data on the non-weighted condition, and this number increased 22.5% to 327 total crossovers in the weighted condition. It also can be clearly seen that there is a exponential decline in crossovers as the targets become farther away into the contralateral space. However, more crossovers can be seen in the weighted condition until the stimuli become so far out that the number over crossovers in total falls below 10 (at −19.2cm). The total distribution of crossovers with respect to contralateral stimulus position can be seen in Fig 6.

**Fig 6.**
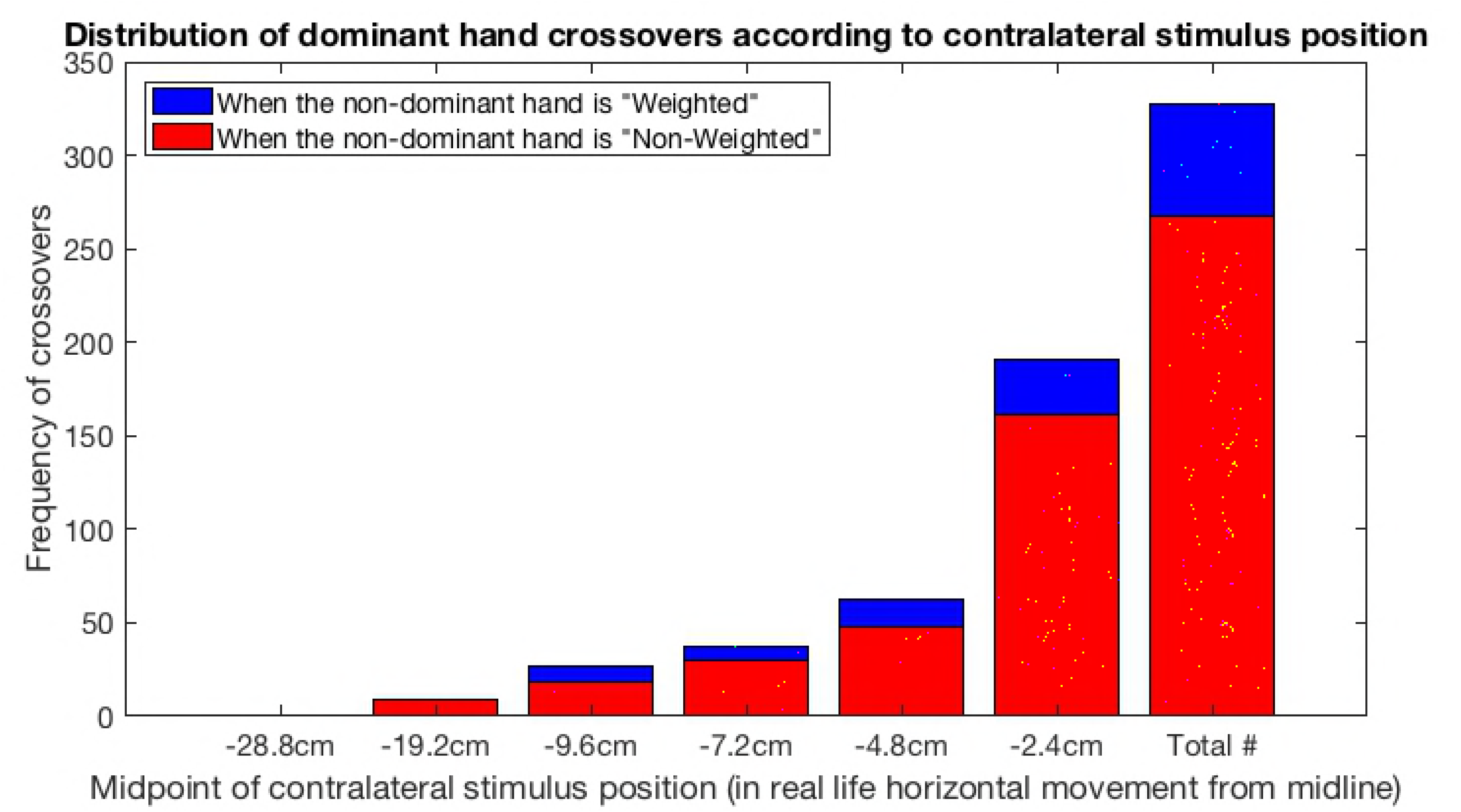
Spatial distributions of dominant (right) hand crossovers. Stimulus positions are listed as accurately as possible as horizontal-plane real-space positions; the red bars represent crossovers in the non-weighted condition, while the blue bars represent the dominant hand crossovers when a weighted glove was on the non-dominant hand.

We asked our participants if they perceived a change in hand preference from the non-weighted to the weighted condition, 16 of the participants reported that they did notice a change in their behavior. There was a space for follow-up information, short answers were given such as: “it was easier to use the non-weighted hand; it was harder to go quickly with the weighted hand; I used the left hand less”. If a participant gave additional input, it can be found in **Supplementary File 2**.

Importantly, our paradigm also showed that we can detect the difference between the weighted and non-weighted conditions even with such a minimal weight. Given we used a 500g weight on the left-hand, it would reason that if we increased this weight, we’d see an even greater increase in right-hand use. Or on the contrary, if the weight was added to the dominant hand and if the *effort-effect hypothesis* should hold, there should be a certain amount of weight relative to each participant that would create roughly a 50%/50% *switch-point*. The rationale behind adding the weighted glove in this paradigm was to make a simple connection to what a patient with an impaired arm may experience; i.e. more difficulty in doing tasks with that limb. We wanted to see if the paradigm was sensitive enough to distinguish the within-subject condition of the added weight. Our findings support that the paradigm indeed is sensitive enough to do so.

Neuropsychologists have a vested interest in understanding hand selection and its relation as a direct link into cerebral lateralization and functioning. Given the validation of our paradigm with the previous literature we believe it can be easily adapted in the future to create a reliable diagnostic instrument and rehabilitative training for patients. For example, after gathering the baseline data of a patient and measuring their *switch-point* for each height level, one would have insight into the extent of the reaching ability compared to controls. If we would see a detriment in left hand use, such as the case in many hemiparesis patients, we could implement a rehabilitative training paradigm. In this next-step adaptation, instead of yellow cubes being presented, we could present ‘red’ or ‘blue’ cubes, corresponding to a ‘red-colored’ left, or a ‘blue-colored’ right hand. Through the rules set by the programming of the environment, the paradigm would only recognize when the red cubes would be contacted by the ‘red hand’, and vice versa. Given our complete control over the stimulus presentation, we could, for example, present these cubes at exactly the same locations as the participant contacted in the baseline paradigm. Yet, we can also challenge the patient to utilize the significantly affected left-hand by programming a training goal to shift the *switch-point* by a certain percentage. Dependent upon the behavior of the patient (i.e., they were successful), the following trials would shift the *switch-point* farther, challenging the patient once again. This sort of adaptive learning paradigm has been shown to be extremely beneficial in rehabilitative efforts, in which the approach is thought to “facilitate motor learning by progressively challenging the subject in accordance with the individual capacity for functional restoration” [37]. Furthermore, rehabilitating left hand use is only one of the many adaptations of the basic paradigm structure we created; easy manipulations can encourage patients to increase accuracy, or speed, or contralateral hand use, or attention and visually guided movements on pre-set trajectories. The virtual environment created here offers a viable approach for flexible rehabilitation paradigms, while its combination with real-time adaptive learning algorithms could be used to tailor therapy for each individual patient.

## Acknowledgments

This work was supported by funds from the Graduate Training Center of Neuroscience, Tuebingen, Germany. We thank Winfried Ilg and Björn Müller, Section for Computational Sensomotorics, Tuebingen for discussions of VR setups and Unity3D programming. We also thank Amin Dadashi for discussions of experimental design and data analysis.

## Supporting information

**S1 File. List of grid positions of cube presentations in Unity3D units.**

[S1 File] Note: the X-axis values are centered on 4.

**S2 File. Questionnaire and responses.**

[S2 File]

